# Comparative scototaxis among four honey bee species

**DOI:** 10.64898/2026.08.18.745666

**Authors:** Yiheng Cai, Fang Liu, Boyan Zhang, Zachary Y. Huang

## Abstract

Honey bees can swim on the water surface toward dark regions, a behavior known as scototaxis that may facilitate escape from water. Although this behavior has been reported in both honey bees and solitary bees, variation among honey bee species remains poorly understood.

We compared scototaxis during swimming in four honey bee species representing two nesting types: open-nesting (*Apis florea* and *A. dorsata*) and cavity-nesting (*A. cerana* and *A. mellifera*). Individual bees were released into a water-filled arena containing a dark sector, and their landing angles were recorded.

All species exhibited significant orientation toward the dark sector. However, open-nesting species showed significantly stronger orientation than cavity-nesting species. No significant differences were detected between replicate colonies within species or between species within the same nesting type, whereas differences between nesting types were highly significant.

Hierarchical clustering based on orientation strength placed *Osmia*, a solitary cavity-nesting bee from our previous study, in the same behavioral cluster as the two open-nesting *Apis* species rather than the cavity-nesting honey bees. We also measured swimming duration, distance, and velocity, but found no consistent differences between nesting types.

These results demonstrate substantial interspecific variation in swimming scototaxis. The behavioral clustering is consistent with the hypothesis that strong scototaxis represents an ancestral trait that has been reduced in the derived *A. cerana*/*A. mellifera* lineage.

## Introduction

Previous studies have shown that honey bees (*Apis mellifera*) can swim on the water surface (Roh & Gharib 2019) and preferentially move toward dark regions (Liu et al. 2026), a behavior known as scototaxis (Fraenkel & Gunn 1940). This behavior is considered adaptive because it may facilitate escape from water by guiding bees toward the shoreline (Liu et al. 2026). Similar swimming behavior has also been observed in solitary bees, suggesting that scototaxis may have originated before the evolution of eusociality (Liu et al. 2026).

Based on our previous finding that the solitary bee *Osmia excavata* exhibits stronger scototaxis than *A. mellifera* (Liu et al. 2026), we hypothesized that scototaxis varies systematically among honey bee species and may be associated with phylogenetic position. To test this hypothesis, we compared scototaxis during swimming in four *Apis* species: *A. florea, A. dorsata, A. cerana*, and *A. mellifera*. These species represent two major nesting types—open nesting and cavity nesting—and occupy distinct positions within the *Apis* phylogeny (Arias & Sheppard 2005; Raffiudin & Crozier 2007; Han et al. 2012).

*Apis florea* and *A. dorsata* are open-nesting species that construct single exposed combs and rely on the sun as a direct reference for waggle-dance orientation (Dyer 2002; Michener 2007). In contrast, *A. cerana* and *A. mellifera* are cavity-nesting species that build nests within enclosed spaces and perform waggle dances relative to gravity in the dark nest environment (von Frisch 1967; Seeley 1995). Phylogenetic analyses consistently place the open-nesting species in more basal positions within the genus *Apis*, whereas the cavity-nesting species form the more derived lineage (Arias & Sheppard 2005; Raffiudin & Crozier 2007; Han et al. 2012). These ecological and evolutionary differences provide a framework for testing whether variation in scototaxis is associated with nesting ecology, phylogenetic position, or both.

In this study, we examined whether the strength of scototaxis differs among honey bee species and whether this variation is associated with phylogenetic position, nesting ecology, or both. In addition, we quantified swimming performance (distance, duration, and velocity) to determine whether interspecific variation in scototaxis is associated with locomotor ability.

## Materials and Methods

### Bee collection and handling

Honey bees were collected from a field site in southwestern China (22.085216° N, 100.891885° E), where all four study species naturally occur. Collections were conducted between 20–22 July and 10–11 September 2025. Worker bees were collected during daytime foraging activity. For *Apis florea* and *A. dorsata*, foragers were collected near their nests using a handheld sweep net. No vacuum devices or chemical immobilization were used. For *A. cerana* and *A. mellifera*, returning foragers were collected inside dummy hives after their original colonies had been relocated a short distance away.

After capture, bees were immediately transferred into ventilated plastic containers (20–30 individuals per container). All bees were provided with a 20% honey solution for approximately 12 h before testing. Individual bees were released into the experimental arena without anesthesia or restraint, and each bee was tested only once.

### Experimental setup

The swimming assay followed Liu et al. (2026) with minor modifications. A transparent glass bowl filled with water served as the swimming arena. The bowl had a diameter of 30 cm at the water surface (circumference 94.75 cm) and a water depth of 10 cm.

A circular sheet of white paper (20 cm high) surrounded the bowl. A single black sector, spanning 72°, provided a dark visual cue.

Experiments were conducted under uniform indoor lighting at 26°C. Illumination was provided by overhead lights (~2,000 lux), with no directional light sources. The arena was arranged to ensure even illumination and minimize shadows and glare on the water surface.

To control for potential directional bias, the position of the black sector was rotated among four orientations relative to the observer (up, right, down, and left). Approximately 25% of individuals were tested under each orientation. Bees were released individually into the center of the bowl while the observer remained stationary. Swimming behavior was recorded using a smartphone camera (60 frames s^−1^; resolution 3648 × 2736 pixels) positioned 60 cm above the arena.

### Measurement of orientation

A frame was extracted from each video at the moment the bee contacted the glass wall. Landing angles were measured from these images using ImageJ (version 1.54g).

Angles were defined as the clockwise angle between the left boundary of the black sector and the landing position of the bee. When bees landed outside the black sector on its left side, angles exceeded 180° and were calculated as 360° minus the angle measured by ImageJ. This procedure standardized angle measurements across trials with different sector orientations.

Twenty bees were randomly selected from each species to measure swimming duration, distance, and velocity using EthoVision XT 17 (Noldus, USA).

### Statistical analyses

To test for dark preference within each trial, chi-square goodness-of-fit tests were used to compare the observed number of bees landing in the dark sector with the expected number under a random distribution. The expected proportion was based on the angular width of the dark sector (72° of 360°, i.e. 20%). To compare the proportions of bees landing in the dark sector between groups, chi-square tests of independence were performed using 2 × 2 contingency tables.

Comparisons were conducted between replicate colonies within species, between species within nesting types, and between pooled nesting types (open-nesting vs. cavity-nesting). For pooled analyses, data from replicate colonies were combined within each group prior to statistical testing. All tests were two-tailed with a significance threshold of *P* < 0.05.

Directional bias in swimming orientation was assessed using the Rayleigh test for circular uniformity. Rayleigh tests were applied to landing angles using the function rayleigh.test() in the R package circular (R version 4.5.2). Significant Rayleigh tests indicate non-uniform orientation and clustering around a common mean direction. Circular histograms (rose plots) were generated using the same package.

Accepted phylogenetic relationships among the studied species were redrawn from published phylogenetic analyses of *Apis* (Arias & Sheppard 2005; Raffiudin & Crozier 2007; Han et al. 2012). Behavioral similarity was assessed by hierarchical clustering (UPGMA) using Euclidean distances calculated from the mean orientation strength (*r*) of each species. For *Apis*, mean *r* values were calculated by averaging the two replicate colonies for each species. Because *Osmia excavata* was previously examined using the same swimming assay and analytical procedures (Liu et al. 2026), its published mean orientation strength (*r*) was included for comparison.

To test whether swimming performance differed among species, we used one-way analysis of variance (ANOVA) with species as a fixed factor. To evaluate differences between nesting types while accounting for species-level variation, we fitted linear mixed-effects models with nesting type (open-nesting vs. cavity-nesting) as a fixed effect and species as a random effect using the lmer() function in the lme4 package (R version 4.5.2; R Core Team 2025).

## Results

Within each species, swimming behavior showed consistent directional patterns across replicate colonies. Circular histograms (rose plots) indicated that all four honey bee species exhibited significant clustering toward the dark sector (Fig. 1A), which was confirmed by highly significant Rayleigh tests for circular uniformity (Supplementary Table 1). The two replicates within each species did not show significant differences (Fig. 1S).

**Figure 1.**
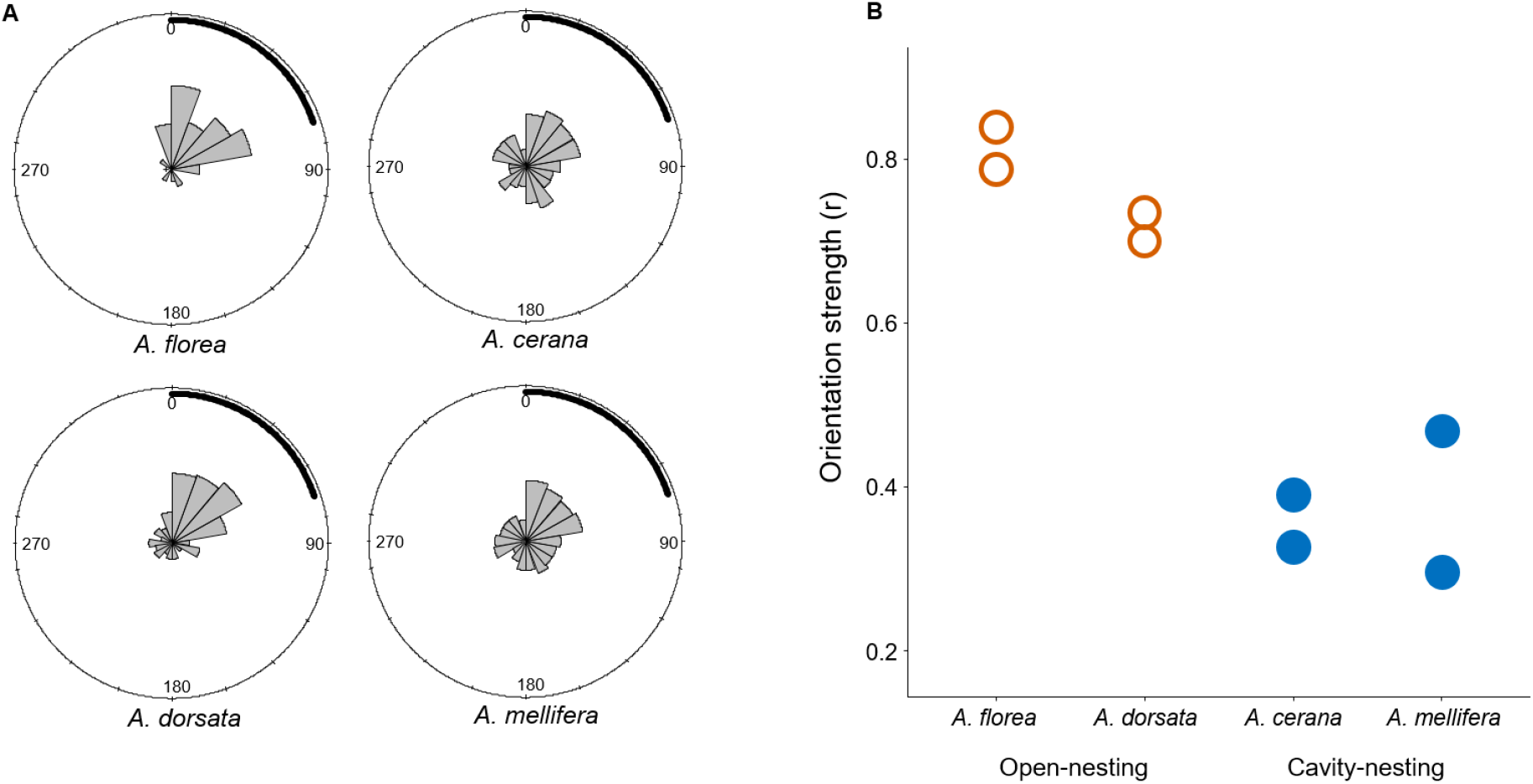
Orientation behavior during swimming in honey bees. **(A)** Circular histograms showing the distribution of individual swimming orientations for each species (*Apis florea, A. dorsata, A. cerana*, and *A. mellifera*). Angles represent the direction of movement relative to the dark area (0 to 72 degrees, inner dark bar), with sectors indicating the frequency of observations. **(B)** Orientation strength (mean resultant length, *r*) for each species, with 2 replicates per species. Higher values of *r* indicate stronger clustering of orientations toward a common direction, whereas lower values indicate more dispersed orientation. Orientation strength differed between nesting types (open-nesting vs. cavity-nesting), but did not differ between species within the same nesting group or between replicates of the same species.

Chi-square comparisons revealed no significant differences in the proportion of landings within the dark sector between replicate colonies of the same species or between species within the same nesting type (all *P* > 0.05; Supplementary Table 2). In contrast, pooled analyses detected a significant difference between nesting types, with open-nesting species showing a higher proportion of landings in the dark sector than cavity-nesting species (*P* < 0.05; Supplementary Table 1B). Consistent with these results, open-nesting species also exhibited higher orientation strengths (*r*) than cavity-nesting species (Fig. 1B).

The accepted phylogeny of the four *Apis* species is shown in Fig. 2A. The behavioral similarity dendrogram (Fig. 2B), however, did not mirror nesting type. *Apis cerana* and *A. mellifera* formed a tight cluster with the lowest orientation strengths (*r*), whereas *Osmia*, despite being a cavity-nesting bee, clustered with the more basal *A. dorsata* and *A. florea* because of its similar orientation strength.

**Figure 2.**
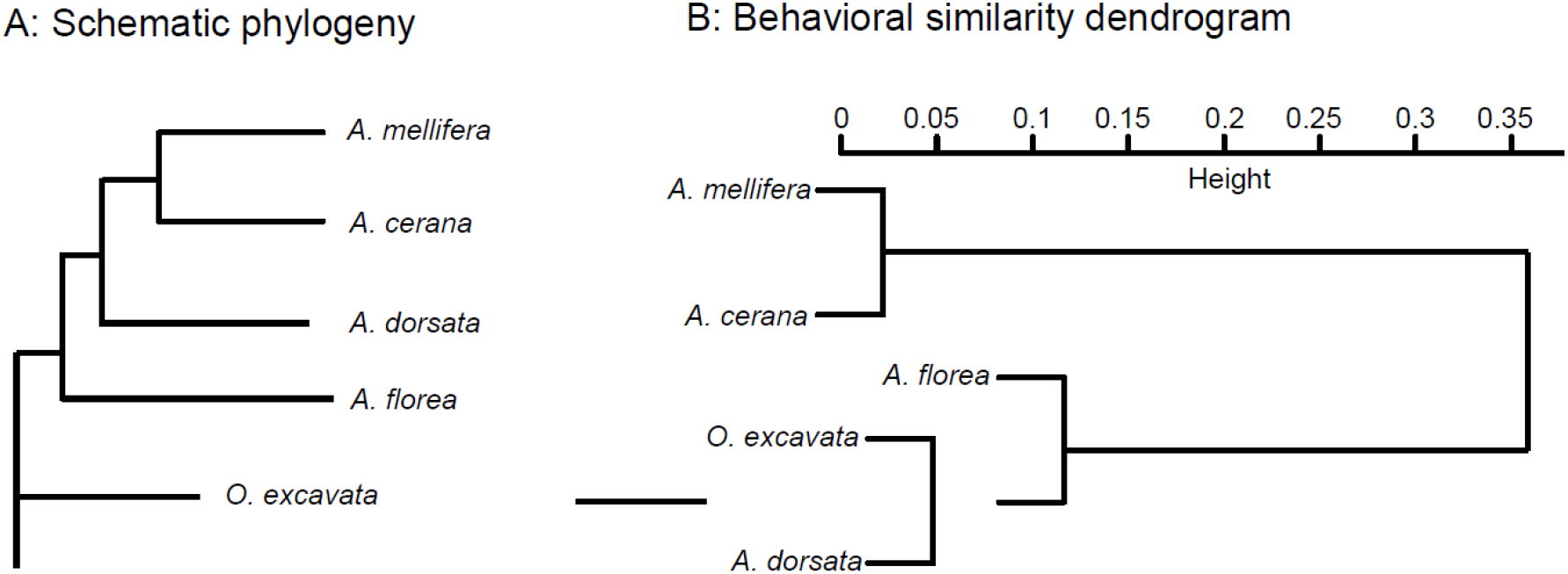
Phylogeny vs swimming behavior. (A) Accepted phylogenetic relationships among the studied taxa, redrawn from published molecular phylogenies (Arias & Sheppard 2005; Raffiudin & Crozier 2007; Han et al. 2012). (B) Hierarchical clustering (UPGMA) of behavioral similarity based on species mean orientation strength (r).

Swimming performance differed among species in total distance (ANOVA: *F*3,76 = 4.10, *P* = 0.009) and showed a marginal difference in duration (*F*3,76 = 2.73, *P* = 0.05), but not in swimming velocity (*F*3,76 = 0.49, *P* = 0.69). Tukey’s HSD tests showed that *A. dorsata* swam significantly shorter distances and for shorter durations than *A. florea* (distance: *P* = 0.005; duration: *P* = 0.028), whereas all other pairwise comparisons were not significant (Fig. 3A). Linear mixed-effects models detected no consistent differences in swimming performance between nesting types.

**Figure 3.**
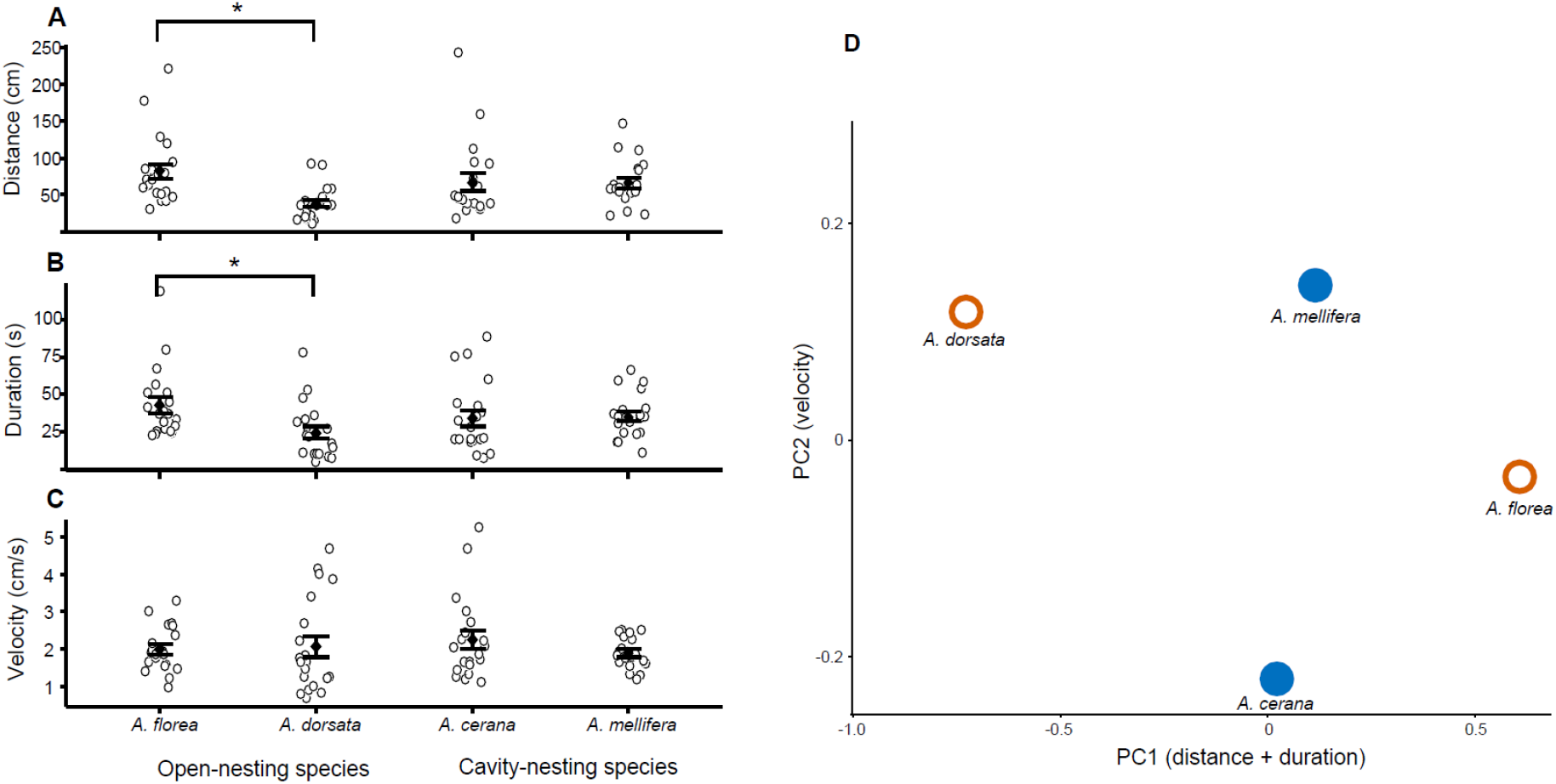
Swimming performance across honey bee species. **(A)** Total swimming distance (cm) for each species. **(B)** Swimming duration (s) for each species. **(C)** Swimming velocity (cm s^−1^) for each species. Each point represents one independent replicate. Asterisks indicate statistically significant differences between groups (see Methods for statistical tests). **(D)** Principal component analysis (PCA) of swimming performance metrics (distance, duration, and velocity). Points represent individual replicates, colored by species. PC1 primarily reflects variation in distance and duration, whereas PC2 reflects variation in velocity.

Principal component analysis (PCA) of swimming performance variables showed that PC1 primarily reflected swimming distance and duration, whereas PC2 was primarily associated with swimming velocity. No clear separation was observed between nesting types or among species of different body sizes (Fig. 3B).

## Discussion

Together, our results reveal a clear pattern: swimming scototaxis is conserved across *Apis*, but its strength differs systematically among species. All four honey bee species showed significant orientation toward the dark sector, indicating that scototaxis is a general water-escape behavior. However, open-nesting species consistently exhibited stronger orientation than cavity-nesting species. This difference was robust across replicate colonies and was not accompanied by consistent differences in swimming distance, duration, or velocity, indicating that variation in scototaxis reflects orientation behavior rather than locomotor performance.

The evolutionary interpretation of this pattern requires caution because nesting ecology and phylogenetic position are confounded within *Apis*. Open-nesting species occupy more basal positions in the phylogeny, whereas cavity-nesting species belong to the derived lineage.

Consequently, the present data alone cannot distinguish whether variation in scototaxis reflects phylogenetic history, nesting ecology, or both. However, when considered together with our previous finding that the solitary cavity-nesting bee *Osmia excavata* exhibits stronger scototaxis than *A. mellifera* (Liu et al. 2026), a different pattern emerges. The behavioral similarity dendrogram grouped *Osmia* with the two open-nesting *Apis* species rather than with the cavity-nesting honey bees. Although this clustering is based on behavioral similarity rather than phylogenetic inference, it is consistent with the hypothesis that strong scototaxis represents an ancestral behavioral trait that is present in *Osmia* and the basal lineages but reduced in the derived *A. cerana*/*A. mellifera* lineage. A more rigorous test of this hypothesis will require broader taxonomic sampling and phylogenetic comparative analyses.

More broadly, these findings add scototaxis to a growing list of behavioral traits that differ between open- and cavity-nesting honey bees, including communication, defense, and activity patterns. Together, these differences suggest that the evolutionary transition from open to cavity nesting was accompanied by coordinated changes in sensory and behavioral systems.

One limitation of this study is that worker caste was not identical among species. *Apis mellifera* and *A. cerana* were sampled as returning foragers, whereas *A. florea* and *A. dorsata* were collected directly from their exposed combs and may have included workers of different ages or task groups. However, if foraging experience enhances visually guided orientation, this sampling difference would be expected to increase, rather than decrease, scototaxis in the cavity-nesting species. Thus, the observed pattern is unlikely to be explained by differences in worker caste, although future studies controlling for age and task would provide a more rigorous test.

Together, these differences suggest that the evolutionary transition associated with cavity nesting may have been accompanied by coordinated changes in sensory and behavioral systems. This variation is independent of swimming performance and, when considered alongside our previous findings in the solitary bee *Osmia excavata*, is consistent with the hypothesis that strong scototaxis represents an ancestral behavioral trait. Broader comparative studies across bee lineages will be needed to test this hypothesis and clarify the relative contributions of phylogenetic history and nesting ecology to the evolution of water-escape behavior.

## Acknowledgement

We thank Guoan Cai and Liu Zhang for financial support (travel and lodging cost for YC to China); Wenfeng Li for teaching YC the swimming assay; Jun Peng, Jingwu Zhang for help in data collection; Yan Wen, and Zhiqiang Qiu for bee collection.

## Author Contributions

YC, data collection, data analysis, first draft; BZ, data collection; FL, swimming performance data analysis, ZYH, experimental design, data analysis and interpretation. All authors participated in writing.

---

**Fig. 1S.**
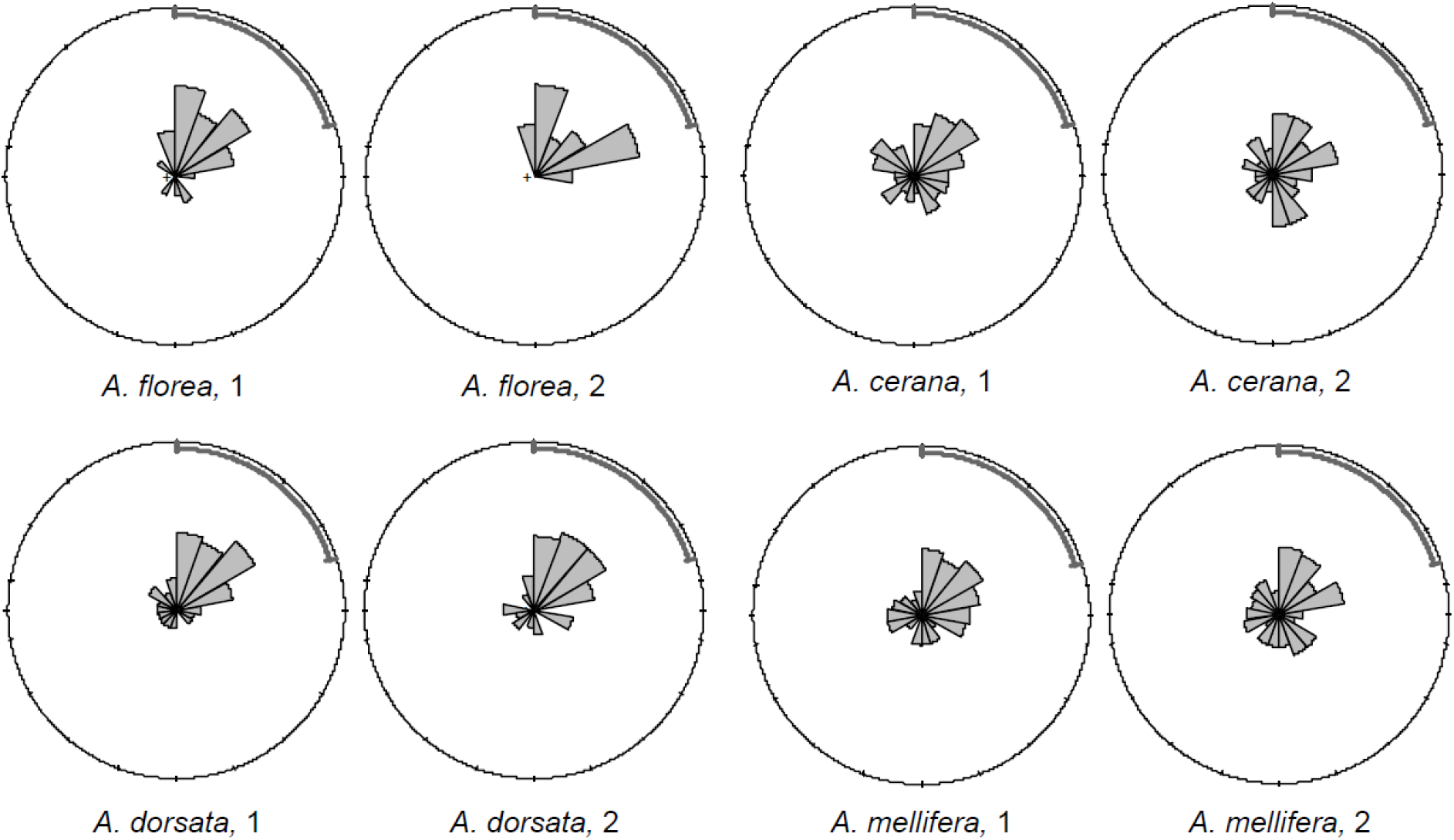
Rose plots of four honey bees species, each with 2 trials. Each trial used a different colony. Contingency Table Analyses indicated no significant differences between each pair of replicate with each species.

**Supplementary Table 1.** Sample sizes and numbers of bees observed in the dark area for each trial. χ^2^ values were calculated using goodness-of-fit tests comparing observed counts with the expected proportion under random distribution (20% of bees in the dark area; null hypothesis: no skototaxis). The *r*value represents the mean resultant length produced by Rayleigh’s test of circular uniformity (rayleigh.test(), *circular* package in R). Each trial used a different colony of bees.

| Species | Trial | Total Bees | Bees in Dark | % in Dark | $\chi^2$ | P value | $r$ | P value |
| --- | --- | --- | --- | --- | --- | --- | --- | --- |
| *A. florea* | 1 | 70 | 57 | 81.4% | 165.09 | <2.2e-16 | 0.799 | 5.73E-19 |
| *A. florea* | 2 | 85 | 66 | 77.6% | 176.54 | <2.2e-16 | 0.851 | 8.16E-26 |
| *A. dorsata* | 1 | 86 | 69 | 80.2% | 195 | <2.2e-16 | 0.745 | 2.35E-20 |
| *A. dorsata* | 2 | 93 | 74 | 79.6% | 206.26 | <2.2e-16 | 0.720 | 1.25E-20 |
| *A. cerana* | 1 | 86 | 45 | 47.7% | 56.17 | 6.66E-14 | 0.411 | 4.87E-07 |
| *A. cerana* | 2 | 92 | 44 | 52.2% | 44.52 | 2.52E-11 | 0.345 | 1.35E-05 |
| *A. mellifera* | 1 | 98 | 55 | 56.1% | 79.92 | <2.2e-16 | 0.484 | 4.12E-11 |
| *A. mellifera* | 2 | 108 | 54 | 50.0% | 60.75 | 6.48E-15 | 0.315 | 1.77E-05 |

**Supplementary Table 2.** Chi-square comparisons within and among species and between the two nesting types. Comparisons were based on the data presented in Supplementary Table 1, using the numbers of bees observed in the dark area and outside the dark area as inputs for contingency table analyses.

| Comparison | $\chi^2$ | *P* value | Significant |
| --- | --- | --- | --- |
| *A. cerana*<br>(between 2 colonies) | 0.36 | 0.549 | No |
| *A. dorsata*<br>(between 2 colonies) | 0.012 | 0.912 | No |
| *A. florea*<br>(between 2 colonies) | 0.34 | 0.563 | No |
| *A. mellifera*<br>(between 2 colonies) | 0.77 | 0.379 | No |
| *A. cerana* vs *A. mellifera* | 0.32 | 0.57 | No |
| *A. dorsata* vs *A. florea* | 0.015 | 0.90 | No |
| Open- vs cavity-nesting | 60.38 | **$7.8 \times 10^{-15}$** | **Yes** |

